# Engineering High Throughput Screening Platforms of Cervical Cancer

**DOI:** 10.1101/2022.10.16.512447

**Authors:** Ines A Cadena, Mina R Buchanan, Conor G Harris, Molly A Jenne, Willie E Rochefort, Dylan Nelson, Kaitlin C Fogg

## Abstract

There is a critical need for complex multicellular three-dimensional physiomimetic models of cancer that can interface with high throughput drug screening methods to assess anti-metastatic and anti-angiogenic drug efficacy in a rapid yet high content manner. We report a multilayer multicellular platform of human cervical cancer cell lines and primary human microvascular endothelial cells that incorporates critical biophysical and extracellular matrix cues, interfaces with standard high throughput drug screening methods, and can evaluate cervical cancer invasion and endothelial microvessel formation over time. Through the use of Design of Experiments statistical optimization, we identified the specific concentrations of collagen I, fibrinogen, fibronectin, GelMA, and PEGDA in each hydrogel layer that maximized cervical cancer invasion and endothelial microvessel length simultaneously. We then validated the optimized platform and assessed the viscoelastic properties of the composite hydrogels as well as their individual constituents. Finally, using this optimized platform, we conducted a targeted drug screen of four clinically relevant drugs on two cervical cancer cell lines. From these data we identified each of the cervical cancer cell lines (SiHa and Ca Ski) as either responsive or refractive to Paclitaxel, Dasitinib, Dovitinib, or Pazopanib. Overall, we developed a phenotypic drug screening platform of cervical cancer that captures cell behavior present in the cervical cancer tumor microenvironment, captures patient to patient variability, and integrates with standard high throughput high content drug screening methods. This work provides a valuable platform that can be used to screen large compound libraries for mechanistic studies, drug discovery, and precision oncology for cervical cancer patients.

## 1. Introduction

Cervical cancer is the second leading cause of cancer-related death in women under 40 and is one of the few cancers to have an increased incidence rate and decreased survival rate over the last ten years.^1^ While HPV vaccination is an effective preventative measure, less than 50% of patients with uteruses aged 13-17 in the United States are up to date with HPV vaccination.^2,3^ Surgery or radiation with chemotherapy are effective first-line treatment strategies for early-stage or locally advanced cervical cancer. However, one in five patients will have recurrent and/or distant metastatic disease, and these patients face a 5-year survival rate of less than 17%, leading to the annual mortality of 4,290 people in the United States alone.^3,4^ The most common clinical treatment for advanced or recurrent cervical cancer is chemotherapy in combination with Bevacizumab, an antibody against vascular endothelial growth factor, to prevent the formation of new blood vessels.^4^ However, while this treatment highlights the importance of blood vessel formation in cervical cancer progression, it only improves survival by 3-4 months, resulting in a patient population that is still greatly underserved.^4^ Thus, there remains a pressing need for novel anti-cancer drugs. However, developing new anti-cancer drugs remains a challenge, as only 7% of novel anti-cancer drugs are approved for clinical use.^5^

Historically, the effectiveness of anti-cancer drugs has been studied *in vitro* using a monolayer of cells seeded on tissue culture plastic. Yet these two-dimensional (2D) monolayer models cannot capture biophysical cues in the tumor microenvironment, such as cell-cell and cell-matrix interactions.^6,7^ Furthermore, cancer metastasis is driven by cancer invasion and angiogenesis which are inherently three-dimensional (3D) processes.^8^ This represents a critical technology gap for evaluating anti-cancer drugs, as compounds targeting angiogenesis and invasion would have immediate clinical relevance.^8^ Thus, there is a growing appreciation for complex multicellular 3D tissue engineered constructs for cancer research and drug discovery.^9–12^ Compared to 2D cell culture, 3D constructs better replicate the tumor microenvironment, resulting in more physiologically relevant responses to chemotherapeutic agents.^6,8,13^ Incorporating 3D constructs into high-throughput drug screening assays is still a developing field, but recent studies have demonstrated its clinical translation potential.^14–17^ However, many 3D assays don’t interface well with standard high throughput screening methods. While most 3D constructs are made in 24-well plates or larger, 96-well plates are the most prevalent format for screening compounds in a high throughput manner.^17^ Furthermore, molecular target based drug screens are classically performed in 2D due to their ease of rapid assessment both through 2D image analysis and the use of high throughput high content multimode plate readers. However, these high throughput assays are difficult to translate to 3D, as confocal imaging is the standard imaging technique for cells in 3D yet is highly time intensive.^18^ Thus, there is a need for 3D tissue engineered tumor microenvironments that can interface with high throughput drug screening methods and can assess anti-cancer drug efficacy in a rapid high content manner. To bridge this gap, there has been a resurgence in phenotypic drug discovery.^17,19^ Instead of evaluating whether a pre-specified molecular target has been modulated, phenotypic drug screening focuses on modulation of cell phenotype from diseased to healthy.^20^ However, to harness the promise of phenotypic drug screening, *in vitro* models of disease must capture biophysical properties of the diseased microenvironment.

The cervical cancer tumor microenvironment is comprised of multiple different extracellular matrix (ECM) structural proteins, soluble factors, and cells.^21^ These elements act in concert to drive cervical cancer invasion and microvessel formation.^22^ While there are limited 3D *in vitro* models of cervical cancer, they have demonstrated increased viability, proliferation, and chemoresistance compared to 2D cultures. However, these models consisted of either a collagen or gelatin hydrogel with cervical cancer cells alone and there are currently no 3D *in vitro* models that can evaluate cervical cancer invasion and microvessel formation simultaneously.^23–26^ Thus, more complex models are needed to better capture the ECM, cell to cell interactions, and cell phenotypic behavior found in the cervical cancer tumor microenvironment.

The overall objective of this study was to develop and validate a 3D *in vitro* model that approximates the cervical cancer tumor microenvironment, interfaces with high throughput drug screening methods, and can evaluate cancer invasion and endothelial microvessel formation over time. We engineered a 3D multilayer multicellular *in vitro* model of cervical cancer in a 96-well plate format that achieves significantly greater cervical cancer growth, cervical cancer invasion, and microvessel length compared to Matrigel, the current gold standard for preclinical cancer drug screening platforms.^12,27,28^ Design of experiments multivariable analysis was used to identify the hydrogel formulations that maximized cervical cancer invasion, cervical cancer growth, endothelial microvessel formation, and endothelial cell growth simultaneously. Then, using the optimized construct, we performed a targeted screen of Paclitaxel, Dovitinib, Pazopanib, and Dasatinib, four clinically relevant therapies.^29^ Overall, this study demonstrates that we developed a phenotypic drug screening platform of cervical cancer that captures cell behavior present in the cervical cancer tumor microenvironment and integrates with standard drug screening methods.

## 2. Materials and Methods

### 2.1. Cell lines and reagents

Unless stated, all reagents were purchased from ThermoFisher (Waltham, MA). Human microvascular endothelial cells (hMVEC) were purchased from Lonza (hMVEC 33226, Walkersville, MD) and used without additional characterization. Cells were expanded in EGM-2 MV media (EBM-2 supplemented with Lonza’s SingleQuot supplements: hydrocortisone, human basic fibroblast growth factor (FGF2), human vascular endothelial growth factor (VEGF), human insulin-like growth factor (IGF), human epidermal growth factor (EGF), ascorbic acid, and gentamycin) and further supplemented with 5% fetal bovine serum (FBS) until used at passage 5. Human cervical cancer cell lines SiHa (ATCC® HTB-35™) and Ca Ski (ATCC CRM-CRL-1550) were purchased from ATCC (Manassas, VA) and used without additional characterization. SiHa were expanded in Eagle’s Minimum Essential Medium (EMEM, ATCC), supplemented with 10% FBS (Sigma-Aldrich, St. Louis, MO) and 1% Penicillin-Streptomycin (P/S, Sigma-Aldrich) until used at passage 5. Ca Ski were expanded in RPMI-1640 Medium (RPMI, ATCC 30-2001) supplemented with 10% FBS and 1% P/S. All cell types were expanded in standard cell culture conditions (37°C, 21% O_2_, 5% CO_2_) and subcultured before they reached 80% confluency.

### 2.2. Hydrogel Fabrication

To overcome the barrier of the hydrogels forming a meniscus in 96-well plates due to capillary action in the small wells, all hydrogels were prepared in specialized clear bottom black-walled 96-well plates that enable cylindrical hydrogels of 10 μL (μ-Plate Angiogenesis 96 Well, IBIDI, Fitchburg, WI). Collagen type I (Col1) hydrogels were prepared by combining high concentration bovine Col1 (~10 mg/mL, FibriCol, Advanced Biomatrix, San Diego, CA), 0.1 □N NaOH, and either 1X or 10X PBS for a final concentration of 0.5, 2.25, or 4 mg/mL Col1, 1X PBS, at pH 7.4. Fibrin hydrogels were prepared by combining fibrinogen from human plasma (30 mg/mL Sigma-Aldrich) with thrombin from human plasma (1000 U/mL, Sigma-Aldrich) in PBS for a final concentration of 50 U/mL thrombin and 0.5, 2.25, or 4 mg/mL fibrinogen. Polyethylene glycol diacrylate (PEGDA, 10% w/v, Advanced BioMatrix) and gelatin methacryloyl (GelMA, 7% w/v, Advanced BioMatrix) were crosslinked using the UV photo crosslinker irgacure 2959 for 1 minute under a 365 nm UV light crosslinking source according to protocols specified by the manufacturer. Growth factor deficient (GFD) Matrigel basement membrane matrix (9.2 mg/mL protein concentration, Corning, MA, USA) was used as a control and prepared according to protocols specified by the manufacturer. All hydrogels were allowed to polymerize for 1 hour in the incubator at 37°C, then rinsed with PBS before cells were seeded on top.

To fabricate composite hydrogels, stock solutions of PEGDA (20% w/v) or GelMA (10% w/v) were cross-linked with stock solutions of Col1 (10 mg/mL), fibrinogen (30mg/mL), or fibronectin (1 mg/mL) in a 1:1 ratio with the ECM components and the pH was adjusted by adding 0.1□N NaOH and either 1X or 10X PBS for a final concentration of 10% w/v PEGDA, 7% w/v GelMA, 0.5, 2.25, or 4 mg/mL Col1, 0.5, 2.25, or 4 mg/mL fibrinogen, and 0.125, 0.175 and 0.25 mg/mL fibronectin. All hydrogels were UV crosslinked and incubated at 37°C for one hour, as described above.

After the hydrogels had formed, cancer or endothelial cells were seeded on top. CellTracker Green (CellTracker™ Green CMFDA Dye) was used to label hMVECs, and CellTracker Red (CellTracker™ Red CMTPX Dye) was used to label the cervical cancer cell lines SiHa and Ca Ski. CellTracker Green-labeled hMVEC were seeded on top of the hydrogels at 20,000 cells in 40 μL of EGM-2 MV per well, and CellTracker Red-labeled SiHa or Ca Ski were seeded on top of the hydrogels at 10,000 cells in 25 μL EMEM per well. The media was changed every 12 hours, and the plates were incubated at 37 °C and 5% CO_2_ in a BioSpa live cell analysis system (Agilent Technologies, Santa Clara, CA) for 45 hours.

### 2.3. Multilayer Hydrogel Fabrication

Multilayer multicellular constructs were designed using a similar method to the single-layer hydrogels described above. To prepare the bottom hydrogel, stock solutions of high concentration bovine Col1 (~10 mg/mL) and fibrinogen (30 mg/mL) were combined with either PEGDA (11.7% w/v) or GelMA (8.7% w/v). The pH was adjusted by adding 0.1□N NaOH and either 1X or 10X PBS for a final concentration of 0.5, 2.5, or 4 mg/mL Col1, 0.5, 2.5, or 4 mg/mL fibrinogen, 10% w/v PEGDA or 7% w/v GelMA, 1X PBS, at pH 7.4. The final gel formulation was pipetted into the IBIDI angiogenesis plates at 10 μL per well. The constructs were UV photocured for 1 minute in a CL-1000 UVP crosslinker (170,000 μJ/cm^2^*min) (UVP, Upland, CA) with the plate placed approximately 6 cm away from the light source. The plate was then incubated for one hour at 37°C with 5% CO_2_ to ensure complete gelation. Endothelial cells were then seeded on top of the bottom hydrogel by pipetting 20,000 CellTracker Green labeled hMVEC cells in 40 μL of EGM-2 MV per well. The endothelial cells were allowed to attach for four hours at 37°C and 5% CO_2_.

To prepare the top hydrogel, stock solutions of high concentration bovine Col1 (~10 mg/mL) and fibronectin (1 mg/mL) were combined with either PEGDA (11.7 %w/v) or GelMA (8.7 % w/v). The pH was adjusted by adding 0.1□N NaOH and either 1X or 10X PBS for a final concentration of 0.5, 2.25, or 4 mg/mL Col1, 0.125, 0.175, or 0.25 mg/mL fibronectin, 10% w/v PEGDA or 7% w/v GelMA, at pH 7.4. The media was then removed from the endothelial cells on top of the bottom hydrogel, and 25 μL of the top hydrogel formulation was pipetted on top of the endothelial cells and crosslinked for 30 seconds as described above. The plate was then incubated for one hour at 37°C with 5% CO_2_ to ensure complete gelation. Finally, cervical cancer cells were seeded on top of the top hydrogel by pipetting 10,000 Cell tracker Red-labeled SiHa or Ca Ski in 10 μL of media per well, with the media comprised of a 1:1 ratio of EGM-2 MV and EMEM (**Fig. 1**). The media was changed every 12 hours, and cells were imaged every 3 hours for 45 hours. As a control, multilayer Matrigel constructs were made using the same methods described above, except the custom hydrogel formulations were replaced with growth factor deficient (GFD) Matrigel in both layers. The media was changed every 12 hours and the plates were incubated at 37 °C and 5% CO_2_ in a BioSpa live cell analysis system for 45 hours.

**Figure 1.**
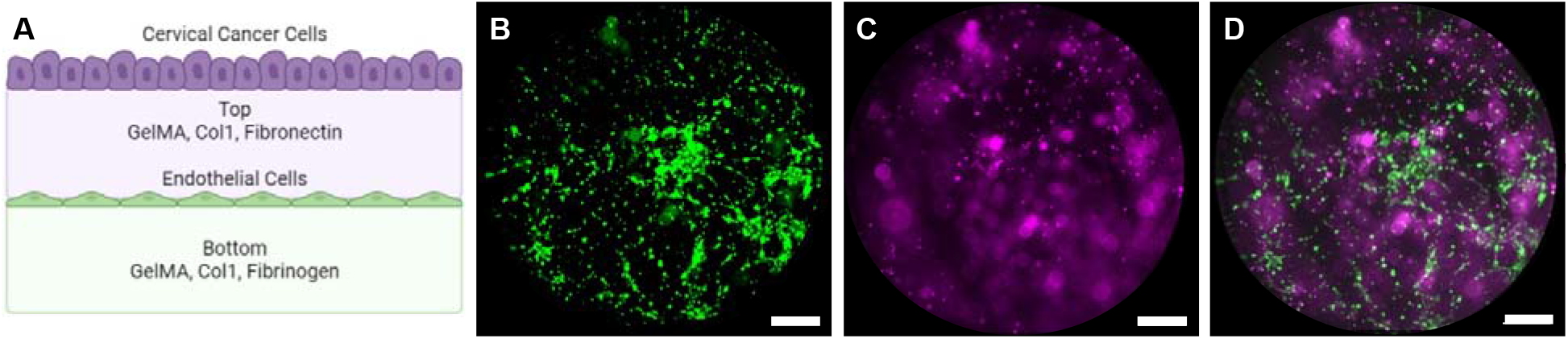
(**A**) Schematic of the multilayer multicellular construct (created with BioRender.com). (**B**) Representative image of CellTracker Green-labeled human microvascular endothelial cells (hMVEC) seeded on the bottom hydrogel. (**C**) Representative image of CellTracker Red-labeled cervical cancer cells seeded on the top hydrogel. (**D**) Merged z-projection image of the endothelial cells and cancer cells in the multilayer multicellular construct. Scale bar represents 1000 μm.

### 2.4. Cellular response to hydrogel formulations

For all experiments, cellular response was evaluated using phenotypic cell metrics. Two-channel 54.8 μm z-stack images were taken every 3 hours for 45 hours after co-culture was initiated using a Cytation 5 cell imaging multimode reader (Agilent Technologies). Images were processed with NIH Fiji-ImageJ and cell coverage was measured for each cell type at every time point by calculating the area within the well covered by cells and dividing it by the total well area. Cell invasion depth was measured using Gen5 software (Agilent Technologies), and endothelial microvessel length was quantified by measuring the average microvessel length at each time point within each well with FIJI (NIH, Bethesda, MD).

### 2.5. Design of Experiments

Design of Experiments statistical optimization was used to identify the significance and interaction of natural and synthetic polymers on cervical cancer and endothelial cell phenotypes. A D-Optimal design was created with MODDE Pro software version 12.1 (Sartorius AG, Göttingen, Germany) to analyze the contribution of four continuous variables (Col1 concentration in the bottom hydrogel, fibrinogen concentration in the bottom hydrogel, Col1 concentration in the top hydrogel, fibronectin concentration in the top hydrogel) and two categorical variables (PEGDA or GelMA in the bottom hydrogel, PEGDA or GelMA in the top hydrogel. The continuous variables were examined at equidistant low, medium, and high levels, resulting in 35 design runs with three centrally repeated conditions to assess the variance in the system (**Supplemental Table 1)**. The ranges for the input variables were as follows: Col1 concentration in the bottom hydrogel ranged from 0.5 mg/mL to 2.5 mg/mL, fibrinogen concentration in the bottom hydrogel ranged from 0.5 mg/mL to 2.5 mg/mL, Col1 concentration in the top hydrogel ranged from 0.5 mg/mL to 2.5 mg/mL, and fibronectin concentration in the top hydrogel ranged from 0.125 mg/mL to 0.175 mg/mL. The output variables were cervical cancer invasion depth, cervical cancer cell coverage, endothelial microvessel length, and endothelial cell coverage. Multiple linear regression was used to generate a predictive model of the output variables as a function of the input variables

### 2.6. Validation of DOE Model

To validate the predictive model, we formulated the hydrogels predicted to either maximize all cell phenotypes measured (cancer invasion, cancer cell coverage, endothelial microvessel length, endothelial cell coverage) or those that would either maximize or minimize cancer invasion or endothelial microvessel length individually. The formulations of these hydrogels were identified using MODDE Pro (**Table 1**).

**Table 1.** Hydrogel formulations predicted to maximize or minimize cell phenotypes of interest in the bottom hydrogel (B) or top hydrogel (T). FG = fibrinogen, Syn = synthetic polymer, FN = fibronectin.

| Response | [Col1]<sub>B</sub><br>(mg/mL) | [FG]<sub>B</sub><br>(mg/mL) | [Syn]<sub>B</sub> | [Col1]<sub>T</sub><br>(mg/mL) | [FN]<sub>T</sub><br>(mg/mL) | [Syn]<sub>T</sub> |
| --- | --- | --- | --- | --- | --- | --- |
| Maximize all cell phenotypes of interest (cancer invasion, endothelial cell microvessel length, cancer cell coverage, endothelial cell coverage) | 2.50 | 2.50 | GelMA | 1.12 | 0.16 | GelMA |
| Maximize cancer invasion | 0.50 | 2.50 | PEGDA | 1.90 | 0.13 | GelMA |
| Maximize endothelial microvessel length | 0.50 | 0.50 | GelMA | 2.50 | 0.15 | PEGDA |
| Minimize cancer invasion | 2.50 | 0.50 | GelMA | 2.50 | 0.14 | PEGDA |
| Minimize endothelial microvessel length | 2.50 | 0.50 | PEGDA | 1.77 | 0.15 | PEGDA |

### 2.7. Assessment of material properties

Dynamic oscillatory shear measurements were used to evaluate the rheological properties of the bottom and top hydrogels of the Maximize all construct as well as GelMA (7% w/v), PEGDA (10% w/v), and GFD Matrigel. The hydrogels with GelMA or PEGDA were photo crosslinked with 365 nm UV light and incubated at 37°C for 1 hour prior to rheological testing. Rheological characterization was conducted using an AR-G2 rheometer (TA Instruments, New Castle, DE) equipped with a 20 mm standard steel parallel top plate with a bottom standard Peltier plate (**Supplemental Fig. 1A**). Due to the tendency for the gels to slip, 150 grit (120 μm particle size) sandpaper was adhered to both the top and bottom fixtures (**Supplemental Fig. 1B**). The gap was adjusted between measurements to ensure contact between the top plate and the gel such that the normal force was 0.1 N. All measurements were performed at a constant temperature of 25 °C using a raised Peltier plate control system. Frequency sweeps were conducted from 0.1 to 100 rad/sec within the linear viscoelastic (LVE) region at a constant strain amplitude of 5%. The storage moduli (*G*’) and loss moduli (*G*”) were measured across the frequency range, the average of the linear region was calculated for each hydrogel, and three consecutive technical replicates were conducted for each hydrogel formulation.

### 2.8. High-throughput high-content drug screening

Using the optimized construct, we performed a targeted drug screen of Paclitaxel, a commonly used chemotherapy agent, and Dovitinib, Pazopanib, and Dasatinib, three clinically used tyrosine kinase inhibitors, which were kindly gifted from the Oregon State University College of Pharmacy High-Throughput Screening Services Laboratory (HTSSL).^29^ The HTSSL provides standard high throughput screening automation equipment, FDA approved drugs, and screening expertise. Cells were seeded in the optimized construct as described above and cultured for 24 hours, at which point they were treated then cultured for another 24 hours. Cells were imaged every 12 hours for the duration of the 48 hour experiment and the response to drug was calculated after 24 hours of treatment. We performed an eight-point half-log screen for all four drugs ranging from 0.008 to 25 μM. Each drug was diluted with dimethyl sulfoxide (DMSO) and pipetted into the wells using an automatic liquid handler (D300e Digital Dispenser, Hewlett Packard (HP), Corvallis, OR). Media alone and DMSO vehicle controls were also evaluated. We then calculated the sensitivity of cervical cancer invasion and endothelial microvessel length in the presence of the four drugs of interest. While IC_50_ values are classically defined as the concentration that inhibits 50% of the cells from proliferating or the concentration that kills 50% of the cells, here we defined the IC_50_ values as the concentration of drug that reduced either cancer invasion, cancer coverage, microvessel length, or endothelial cell coverage to 50% of the value observed in the absence of drug. We calculated the IC_50_ values from the results of the eight-point half-log dose-response studies for both SiHa and Ca Ski cervical cancer cell lines. Specifically, we normalized each cell response to the average of the values observed in the absence of drug and each replicate was considered as an individual point. The IC_50_ and Hill slope was determined from these results (Equation 1).

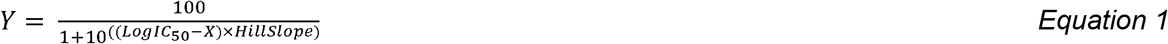

Where, Y is the normalized cell response and X is the log of the drug concentration.

### 2.9. Statistical analysis

Statistical analyses were performed using a two-way analysis of variance (ANOVA) with Tukey correction for multiple comparisons or one-way ANOVA when appropriate. Dunnet correction was used in the DOE comparison to account for the significant difference between the predicted hydrogels and Matrigel. All statistical analyses were performed using Prism 8.2.1 software (GraphPad, San Diego, CA); *p* values less than .05 were considered statistically significant. Replicates are specified in each experiment, and asterisks denote significance.

## 3. Results

### 3.1. Single layer monoculture constructs

While the collagen, fibronectin, fibrinogen, GelMA, and PEGDA have been widely used in biomaterial constructs, the concentration dependent effects of these proteins and polymers on cervical cancer behavior remained unknown. Thus, we investigated the effects of different hydrogel formulations on cervical cancer coverage and invasion, seeding cervical cancer cells alone on top of a single hydrogel layer. We performed an initial screen of GelMA and PEGDA hydrogels, ranging from 5-10% w/v and 10-20% w/v, respectively (**Supplemental Figure 2**). From this screen, we elected to use 7% GelMA and 10% PEGDA, as those concentrations best supported both cervical cancer and endothelial cell behavior. For each type of ECM protein (Col1, fibrinogen, fibronectin) we examined a low, medium, and high concentration with or without a synthetic polymer backbone (**Figure 2**).

**Figure 2.**
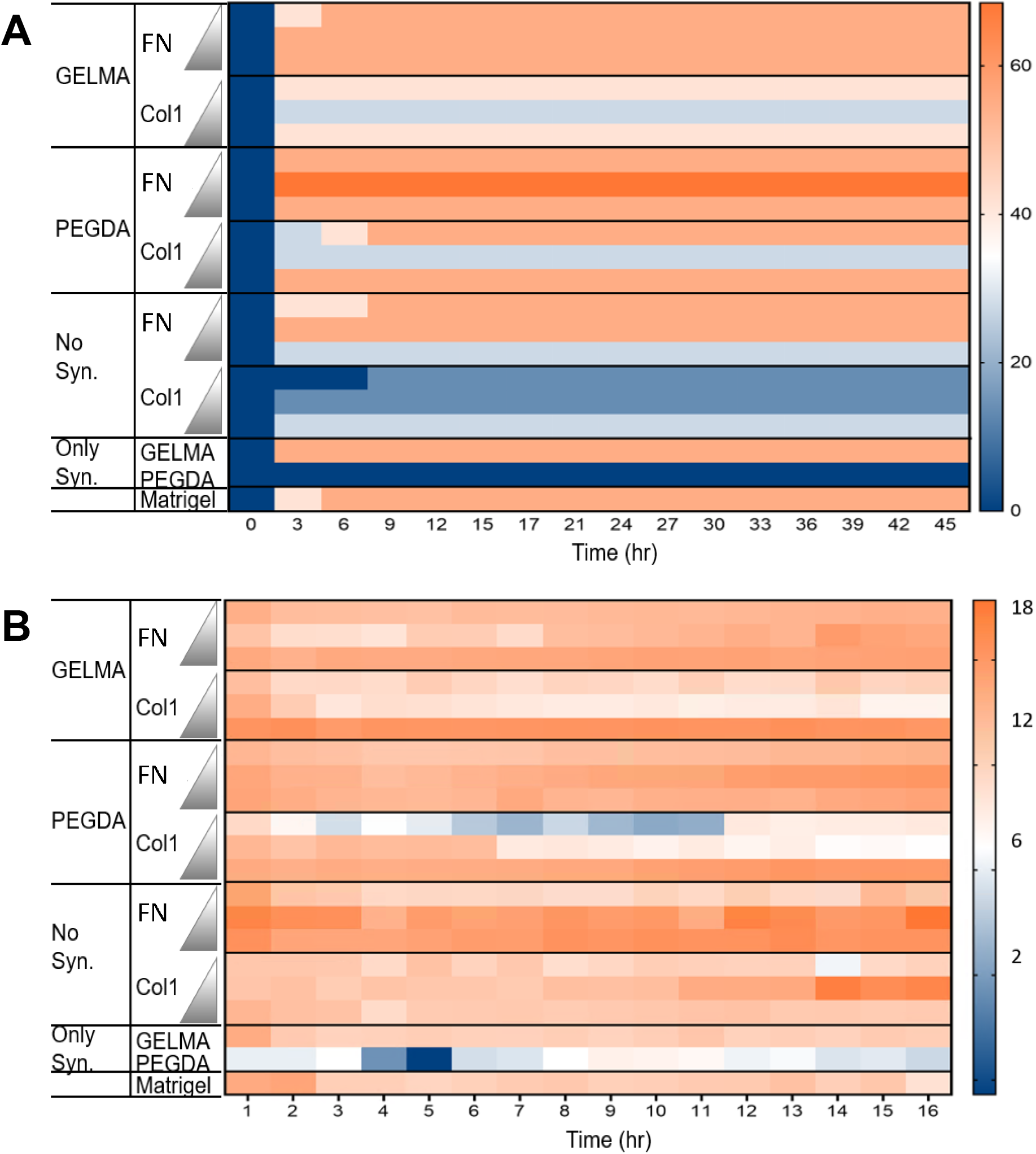
Single layer hydrogels with cervical cancer. (**A**) Cervical cancer cell coverage over time (**B**) Cervical cancer invasion over time. Images were taken and analyzed every three hours for 45 hours. Col1 (0.5, 2.25 and 4 mg/mL) and fibronectin (FN, 0.125, 0.175 and 0.250 mg/mL) were evaluated at low, medium, and high concentrations with 7% w/v GelMA, 10% w/v PEGDA, or alone (No Syn.). 7% GelMA, 10% w/v PEGDA, and Matrigel were evaluated as controls. Triangles represent the lowest to the highest concentration of each natural polymer. Data represented the mean ± SD (n = 4).

Cervical cancer coverage was defined as the percentage of the well covered by cervical cancer cells and cervical cancer invasion was defined as the deepest point that the cervical cancer cells invaded from their initial z position, both of which were quantified every three hours for 48 hours. Collagen, fibronectin, PEGDA, and GelMA were evaluated either alone or in combination with each other, and growth factor deficient (GFD) Matrigel was used as a control. We observed significant differences in cervical cancer coverage (**Figure 2A**) and invasion (**Figure 2B**), depending on the hydrogel formulation that they were seeded on. Specifically, PEGDA alone (10% w/v) did not yield significant changes in coverage from its initial seeding density of 5%. Yet when PEGDA was combined with fibronectin (0.125, 0.175, 0.25 mg/mL) coverage was significantly increased, with the highest coverage observed at a blend of 0.175 mg/mL fibronectin and 10% w/v PEGDA of the fibronectin/PEGDA hydrogels. When PEGDA was combined with Col1 (1, 2.5, 4 mg/mL), coverage increased as Col1 concentration increased, with a blend of 4 mg/mL Col1 and 10% w/v yielding the highest coverage of the Col1/PEGDA hydrogels. Examining GelMA, GelMA alone (7% w/v) had a cell coverage of approximately 7%. Combining GelMA with fibronectin (0.125, 0.175, 0.25 mg/mL) did not yield significant differences in cervical cancer coverage. However, when GelMA was combined with 4 mg/mL Col1, coverage was increased to 15%. Examining the effects of Col1 and fibronectin alone on cervical cancer coverage demonstrated a preference for the middle concentration for both Col1 and fibronectin.

Changing the hydrogel formulation also significantly affected cervical cancer invasion (**Figure 2B**). After 12 hours, most of the invasion occurred for all hydrogel formulations. Cells seeded on top of PEGDA alone did not invade at all. However, PEGDA combined with fibronectin (0.175 mg/mL) had the highest invasion (80 μm). In contrast, cervical cancer cells moderately invaded into GelMA (40 μm). There were no significant differences observed between different collagen concentrations blended with either PEGDA or GelMA.

### 3.2. Developing multilayer multicellular constructs of the cervical cancer tumor microenvironment

To capture the architecture of unique layers of cells and matrix observed histologically in cervical cancer patient biopsies,^30^ we used two distinct hydrogel layers to support the endothelial cells and cervical cancer cells (**Figure 1**). Human microvascular endothelial cells (hMVECs) were used to model cervical vasculature and a human cervical cancer cell line from a primary squamous cell carcinoma (SiHa) was used to model cervical cancer invasion and growth. To interface with standard drug screening methods, we used specialized 96-well plates that enable cylindrical hydrogels of 10 μL with no meniscus formation. Our overall engineering objective was to identify the concentrations of ECM proteins and which synthetic polymer should be used in each hydrogel layer to maximize cell phenotypes that would be present in the cervical cancer tumor microenvironment and advantageous for a novel anti-cancer drug to inhibit. To achieve this objective, we used Design of Experiments (DOE). DOE is a powerful statistical modeling tool that identifies individual and combinatorial effects of multiple input variables on multiple output variables while reducing the total number of experiments required compared to a full factorial approach. While DOE is underutilized in tissue engineering,^31^ it has great potential to engineer biomaterials that maximize or minimize cell phenotypes of interest. For this application, we used DOE to determine the significance and interaction between six variables derived from hydrogel synthesis on four output variables that captured metastatic behavior. Specifically, we analyzed the contribution of four continuous input variables (the concentrations of Col1 and fibrinogen in the bottom hydrogel, the concentrations of Col1 and fibronectin in the top hydrogel) and two categorical input variables (the choice of either GelMA or PEGDA as the synthetic polymer backbone in the bottom and top hydrogels) on four continuous output variables (cervical cancer coverage, cervical cancer invasion, endothelial cell coverage, endothelial microvessel length) (**Table 2**).

**Table 2.** DOE input variables and specific levels tested. B = bottom hydrogel, T = top hydrogel, FG = fibrinogen, Syn = synthetic polymer, FN = fibronectin.

| DOE Input Variable | Variable Abbreviation | Specific Conditions Tested |
| --- | --- | --- |
| [Col1]<sub>B</sub> | A | 0.5, 1.25, 2.5 mg/mL |
| [FG]<sub>B</sub> | B | 0.5, 1.25, 2.5 mg/mL |
| Syn<sub>B</sub> | C | GelMA, PEGDA |
| [Col1]<sub>T</sub> | D | 0.5, 1.25, 2.5 mg/mL |
| [FN]<sub>T</sub> | E | 0.125, 0.150, 0.175 |
| Syn<sub>T</sub> | F | GelMA, PEGDA |

A D-optimal experimental design generated 33 unique construct formulations with a centrally repeating condition (**Supplemental Table 1**). For each condition, we measured the resulting cervical cancer coverage, cervical cancer invasion, endothelial coverage, and endothelial microvessel length at 0, 12, and 24 hours after initiating co-culture (**Figure 3**). For endothelial microvessels, they began to break apart after 12 hours so the values at 24 hours were not easily quantified. For the response surface maps, the cervical cancer coverage at 24 hours, the cervical cancer invasion at 24 hours, the endothelial coverage at 24 hours, and the endothelial microvessel length at 12 hours were used as output variables. A multilayer multicellular construct with GFD Matrigel as the biomaterial in both hydrogel layers was used as a control. All of the output variables measured were significantly influenced by all of the input variables (*p ≤* 0.05) with a coefficient of correlation (R^2^) > 0.8 for all output variables, indicating a good model fit. Additionally, all four output variables had significant interaction terms, which were described as quadratic or two-factor interactions (**Table 3**).

**Figure 3.**
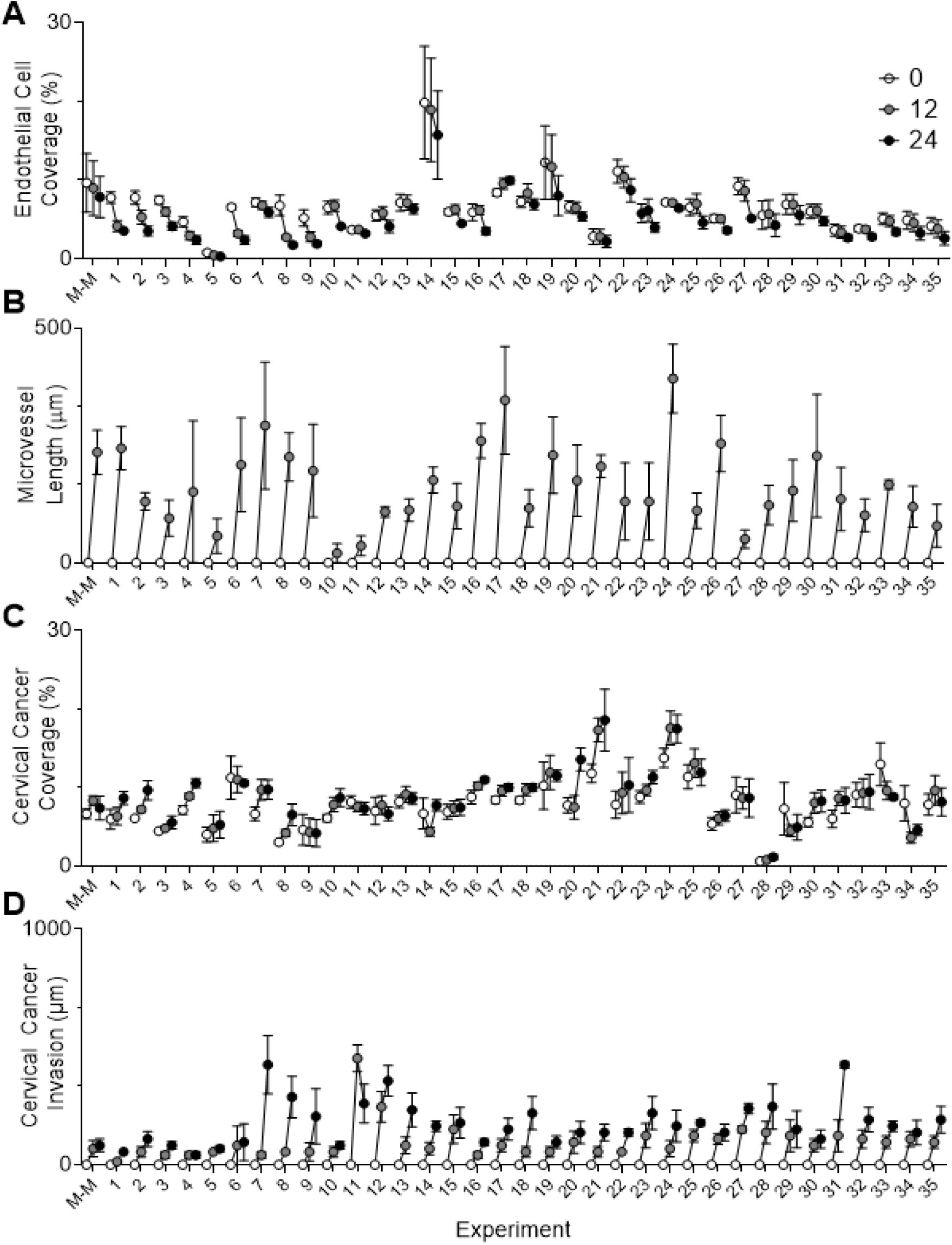
Phenotypic cell responses for each of the 33 unique construct formulations for the D-optimal experimental design with a centrally repeating condition. Images were taken and analyzed every 12 hours for 24 hours. (**A**) endothelial cell coverage, (**B**) microvessel length, (**C**) cervical cancer coverage, (**D**) cervical cancer invasion. Data represented the mean ± SD (n = 4). Matrigel (M-M) was used as a control.

**Table 3.** Summary of DOE model and interplay between input and output variables. Input variables detailed in Table 2. R^2^ = coefficient of correlation.

| DOE Output Variable | $R^2$ | Range Observed | Significant Input Variables<br>$p \leq 0.05$ |
| --- | --- | --- | --- |
| Cervical cancer coverage | 0.87 | 0.4 – 20% | B, C, D, E, F<br>$A^2, B^2$<br>$A^*F, D^*E, D^*B,$ |
| Cervical cancer invasion | 0.94 | 0 – 452 $\mu\text{m}$ | D, F<br>$D^2, E^2$<br>$A^*C, B^*D, B^*F, C^*D^*, C^*F$ |
| Endothelial cell coverage | 0.91 | 0.7 – 19% | A, C, E, F<br>$A^2$<br>$A^*B, A^*C, B^*D, D^*E, E^*C,$ |
| Endothelial microvessel length | 0.80 | 10 – 220 $\mu\text{m}$ | G<br>$A^2, B^2, E^2$<br>$A^*D, B^*D, C^*D$ |

By modulating the concentrations of ECM proteins and the choice of synthetic polymer backbone, we observed cervical cancer coverage ranging from 0.4 to 20% (**Figure 4A, B**). Cervical cancer coverage was significantly influenced by all of the input variables assessed in both hydrogel layers, including the formulation of the bottom hydrogel that did not come in direct contact with the cervical cancer cells upon seeding. Cervical cancer coverage decreased both with increasing fibronectin concentration in the top hydrogel and increasing fibrinogen concentration in the bottom hydrogel. Cervical cancer invasion was also significantly influenced by all of the input variables assessed in both hydrogel layers (**Figure 4C, D**). The interplay of the six input variables yielded cervical cancer invasion ranging from 0.7 μm to 452 μm in depth. For both cervical cancer coverage and cervical cancer invasion, GelMA had a positive influence compared to PEGDA.

**Figure 4.**
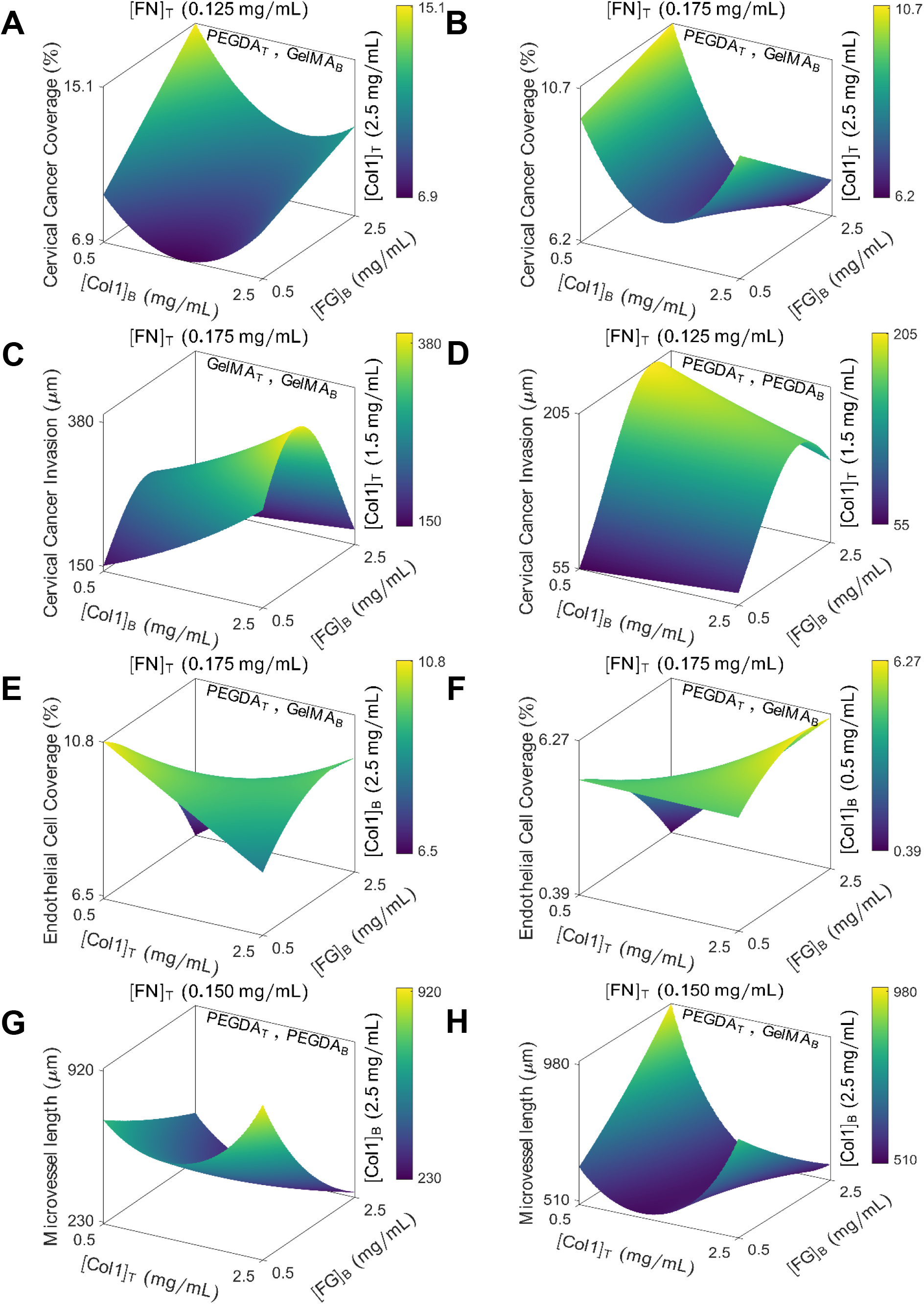
Response surface maps of hydrogel formulation effects on (**A-B**) Cervical cancer coverage (**C-D**) Cervical cancer invasion (**E-F**) Endothelial cell coverage (**F-G)** Microvessel length. Model fit with multiple linear regression

Endothelial cell coverage and microvessel length were also significantly influenced by all of the input variables examined, including the formulation of the top hydrogel that is seeded on top of the endothelial cells. Endothelial cell coverage ranged from 0.7 – 19% and increased quadratically with increasing Col1 concentration in the bottom hydrogel (**Figure 4E, F**). The interplay of the six input variables yielded microvessel lengths ranging from 10 – 220 μm. Microvessel length increased quadratically with increasing with increasing Col1 concentration in the bottom hydrogel yet decreased quadratically with increasing fibronectin concentration in the top hydrogel (**Figure 4G, H**). Taken together, these data indicate that both cervical cancer and endothelial cell behavior were influenced by the ECM proteins and the choice of synthetic polymers in both hydrogel layers.

### 3.3. Validation of DOE Model Results

The desirability function algorithm was used to identify the hydrogel formulations predicted to maximize all four of the cell phenotypic behaviors simultaneously, maximize cervical cancer invasion, maximize cervical cancer coverage, maximize endothelial microvessel length, or maximize endothelial cell coverage. While there were significant differences between groups, cervical cancer and endothelial cell coverage did not exhibit a large dynamic range (**Supplemental Figure 3**). Thus, we chose to focus on cervical cancer invasion and microvessel length moving forward.

The construct that maximized all cell phenotypes (Maximize All) was made with 2.5 mg/mL Col1 and 2.5 mg/mL fibrinogen in the bottom hydrogel, 1.12 mg/mL Col1 and 0.16 mg/mL fibronectin in the top hydrogel, and GelMA was used as the synthetic polymer in both layers. This construct resulted in the highest invasion depth (**Figure 5A**) and microvessel length (**Figure 5B**), a significant increase from all other constructs evaluated. Furthermore, this optimized construct outperformed Matrigel by 10-fold in terms of cervical cancer invasion and 3-fold in terms of microvessel length.

**Figure 5.**
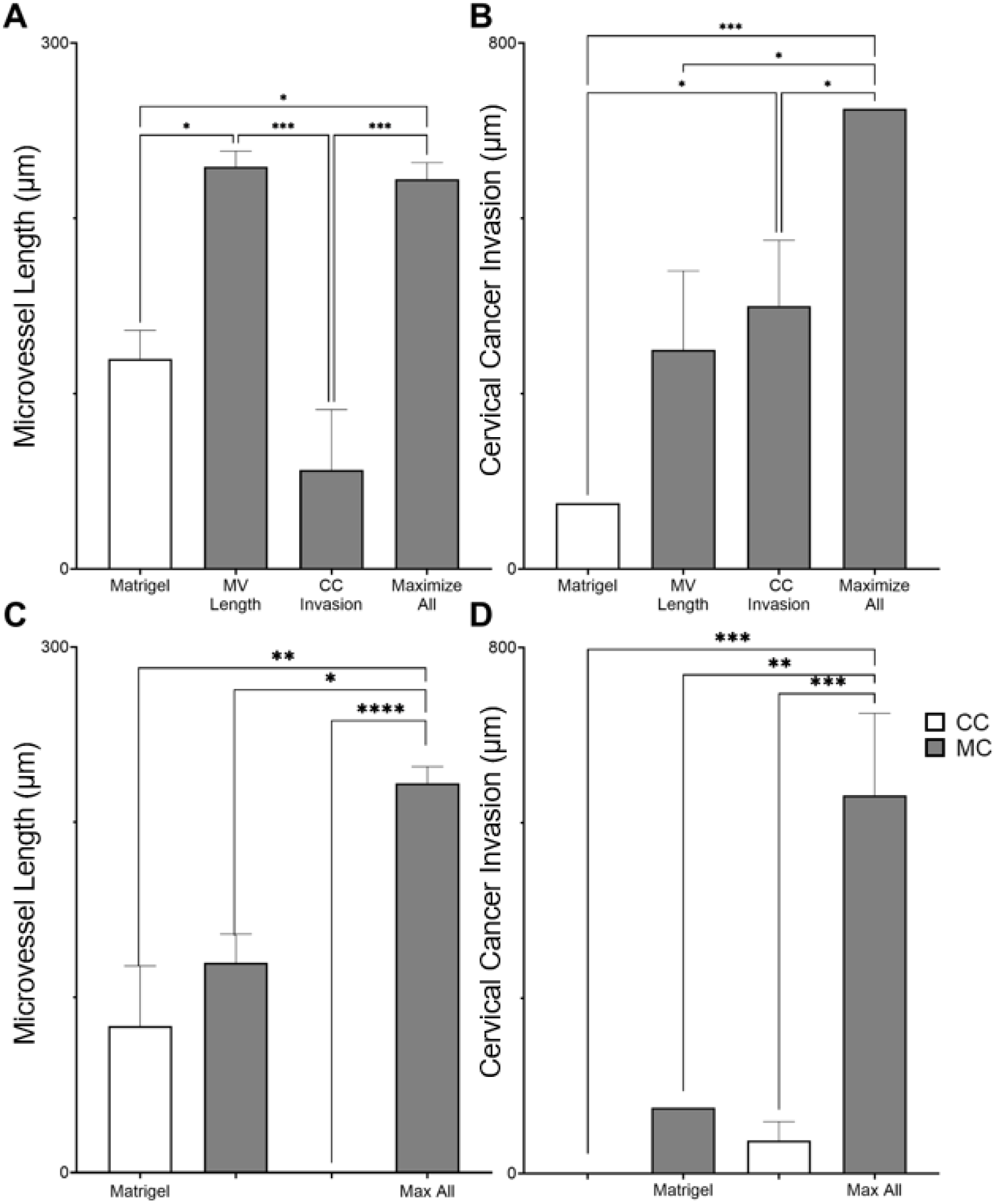
The desirability function algorithm was used to identify the hydrogel formulations predicted to maximize all four of the cell phenotypic behaviors simultaneously (Maximize All), maximize cervical cancer invasion (CC Invasion), or maximize endothelial microvessel length (MV length). **(A)** Microvessel length for optimized hydrogel combinations at 12h. **(B)** Cervical cancer invasion for optimized hydrogel combinations at 48h. To evaluate the contribution of crosstalk between cervical cancer and endothelial cells, each cell type was cultured on the “Maximize all” construct in coculture (CC) or monoculture (MC). **(C)** Microvessel length at 12h **(D)** Cervical cancer invasion at 48 h. Matrigel was used as a control. * *p* <0.03, ** *p*<0.0021, *** *p* <0.0002 compared with the mean of each group. One-way ANOVA with Tukey post-test. Data represent the mean ± SD (n = 4).

To evaluate the contribution of crosstalk between cervical cancer and endothelial cells, each cell type was cultured in the optimized construct with both hydrogel layers present (Maximize all) in the presence or absence of the other cell type and evaluated over time. Again, as a control, the same experiment was performed in a multilayer Matrigel construct. When cervical cancer cells were cultured in monoculture, they did not invade at all on Matrigel and only had slight invasion on the optimized construct (**Figure 5C**). Similarly, when endothelial cells were cultured in monoculture, they did not form any microvessels on the optimized construct (**Figure 5D**). The endothelial cells formed very short microvessels on Matrigel, both in monoculture and in co-culture, with no significant differences observed. Again, for both cervical cancer invasion and microvessel length, the optimized construct drastically outperformed the Matrigel construct. Cervical cancer and endothelial cell coverage were also measured in monoculture and in co-culture, with significantly increased coverage in co-culture compared to monoculture (**Supplemental Figure 4**).

### 3.4. Evaluation of material properties

To evaluate the contribution of mechanical properties on the differences in cellular behavior, the viscoelastic properties of the top and bottom hydrogel layers in the optimized construct were compared to their individual ECM components of fibrinogen and collagen as well as GelMA, PEGDA, and Matrigel (**Figure 6**). Specifically, we assessed the storage modulus (G’) and loss modulus (G”) of the optimized bottom hydrogel (2.5 mg/mL Col1, 2.5 mg/mL fibrinogen, 7% w/v GelMA), the optimized top hydrogel (1.12 mg/mL Col1, 0.16 mg/mL fibronectin, 7% w/v GelMA), the Col1 concentration in the optimized bottom hydrogel (2.5 mg/mL Col1), the fibrinogen concentration in the optimized bottom hydrogel (2.5 mg/mL fibrinogen), the collagen concentration in the optimized top hydrogel (1.12 mg/mL Col1), 7% w/v GelMA, 10% w/v PEGDA, and GFD Matrigel (9.2 mg/mL). All of the hydrogels exhibited viscoelastic behavior, with higher storage moduli than loss moduli and flat plateaus in the linear viscoelastic region, indicating the materials were gels. Those that had larger gaps between the storage and loss moduli, such as GelMA, were more elastic than those that had smaller gaps between the storage and loss moduli, such as 1.12 mg/mL Col1. Of all hydrogels examined, PEGDA had the highest storage modulus followed by GELMA. The synthetic polymers had orders of magnitude higher storage moduli compared to the natural polymers, and as the optimized hydrogels were a combination of the natural and synthetic polymers, their storage moduli fell between the two. For the natural polymers, the 2.5 mg/mL fibrinogen had the highest storage modulus, followed by the 2.5 mg/mL Col1 hydrogel, and lastly the 1.12 mg/mL Col1 hydrogel had the lowest storage modulus of all hydrogels evaluated. Matrigel had a storage modulus higher than any of the other natural polymers, but still significantly lower than the optimized hydrogel layers. Similar observations were made for the loss moduli (G”). Again, PEGDA had the highest loss modulus, and the natural polymers had the lowest loss moduli. There were no statistical differences between GelMA, Matrigel, or the optimized hydrogel layers.

**Figure 6.**
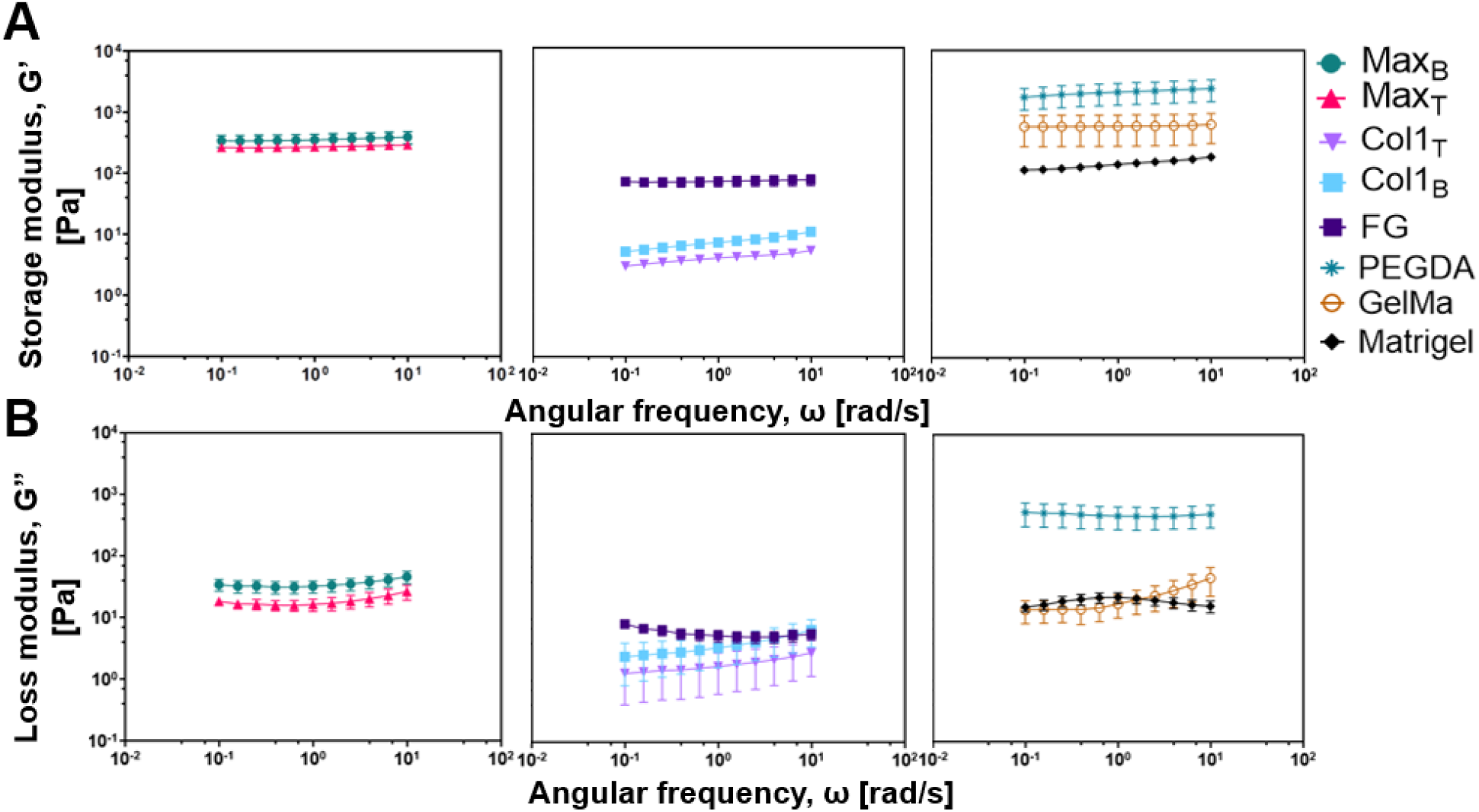
Rheological characterization of hydrogels. Storage moduli (G’) and loss moduli (G”) for the optimized bottom hydrogel (MaxB: 2.5 mg/mL Col1, 2.5 mg/mL fibrinogen, 7% w/v GelMA), the optimized top hydrogel (Max_T_: 1.12 mg/mL Col1, 0.16 mg/mL fibronectin, 7% w/v GelMA), the collagen concentration in the optimized top hydrogel (Col1_T_: 1.12 mg/mL Col1), the Col1 concentration in the optimized bottom hydrogel (Col1B: 2.5 mg/mL Col1), the fibrinogen concentration in the optimized bottom hydrogel (FGB: 2.5 mg/mL fibrinogen), 10% w/v PEGDA, 7% w/v GelMA, and GFD Matrigel (9.2 mg/mL). Frequency sweeps were conducted from 0.1 to 100 rad/sec within the linear viscoelastic (LVE) region at a constant strain amplitude of 5%. Data represent the mean ± SD (n = 3).

### 3.5. High throughput high content phenotypic drug screening

As proof of concept that our constructs can be used as a platform for high throughput high content phenotypic drug screening, we performed an eight-point half-log dose response study of four clinically relevant drugs on two cervical cancer cell lines. We again used SiHa, a human cervical cancer cell line from a primary squamous cell carcinoma, and also evaluated the responsiveness of Ca Ski, a human cervical cancer cell line from an intestinal metastasis of an epidermoid cervical carcinoma. Cervical cancer invasion and endothelial microvessel length were measured as a function of drug concentration (**Figure 7**). The IC_50_ values were calculated for each drug on each cell line, with a higher IC_50_ value indicating drug resistance. The Hill slope was also measured, as it mathematically describes the slope of the dose response curve, with an increased negative Hill slope indicating a stronger drug-receptor interaction.^32^ A positive Hill slope indicates an increased cell phenotypic response with increasing concentration of drug, a hallmark of drug resistance (**Table 5**).^32^

**Figure 7.**
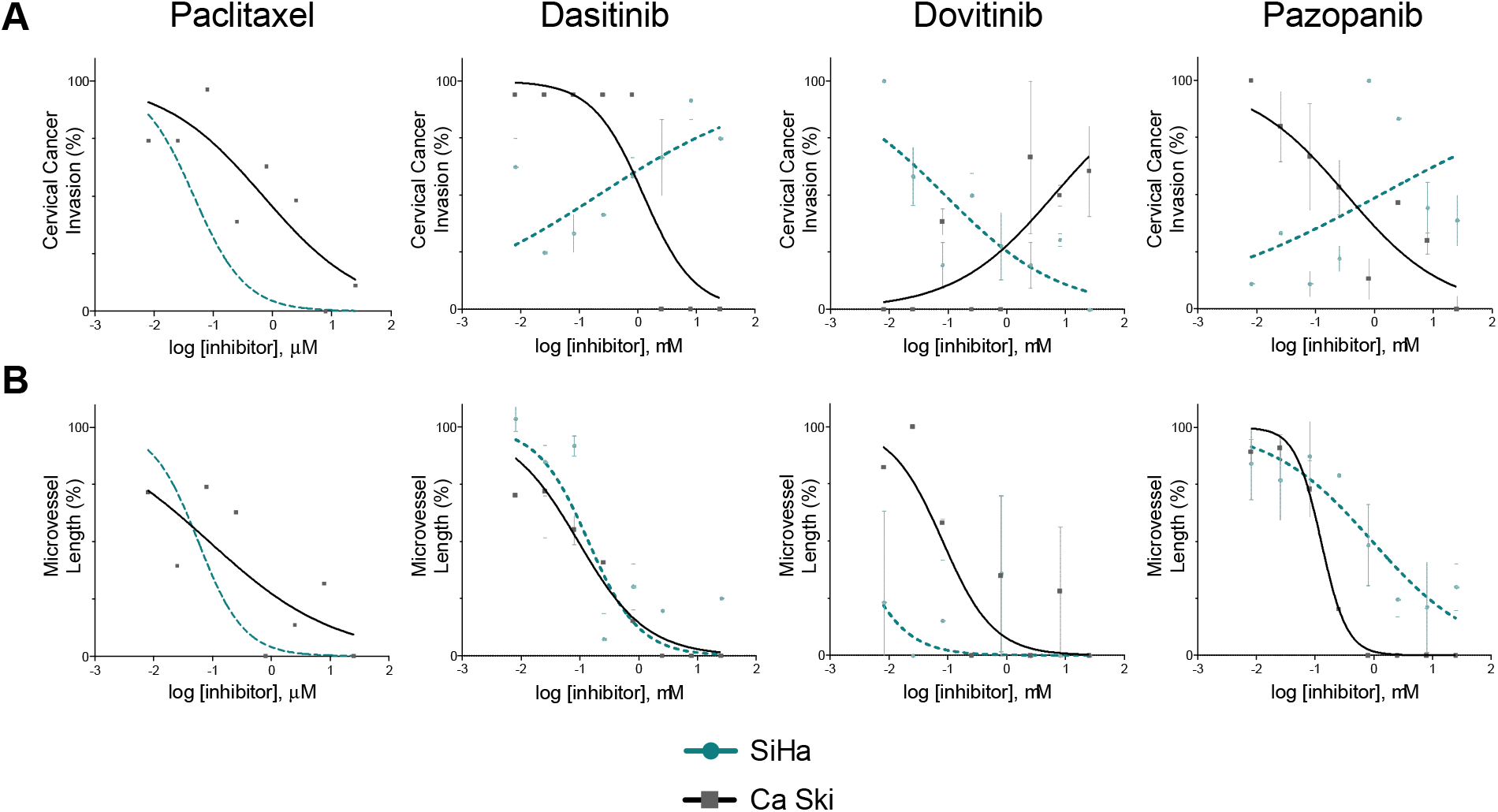
*In vitro* growth-inhibitory effects of the inhibitory drugs on human microvascular endothelial cells (hMVECs) and human cervical cancer cell lines (SiHa, Ca Ski). Cells were seeded in the construct, cultured for 24 hours, then treated with 0.008 – 25 μM of drug for 24 hours, at which point cell response was evaluated. Each cell response is normalized to the average of the values observed in the absence of drug. **(A)** Cervical cancer invasion **(B)** Microvessel length. Data represent the mean ± SD (n = 3).

**Table 4.**
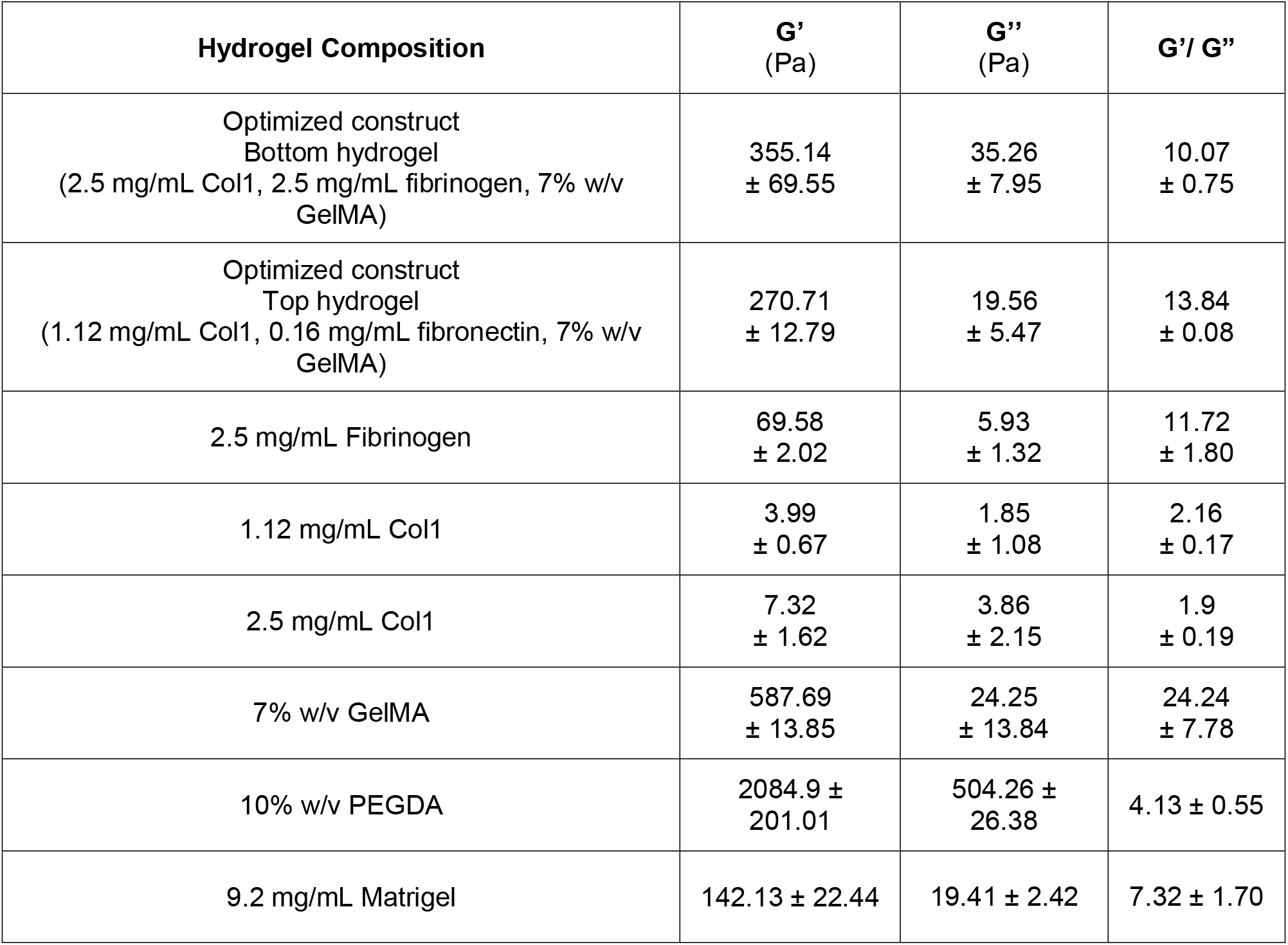
Comparison among the storage moduli (G’), loss moduli (G”) and G’/G” ratio among the different hydrogel combinations. All data = mean ± SD, N=3.

**Table 5.** Comparison of the inhibitory effect of different drugs in multilayer multicellular *in vitro* model for cervical cancer. IC_50_ and Hill Slope values for microvessel length and cervical cancer invasion. Values represent mean ± SEM, N=3.

| Cell line | Inhibitory Agent<br>Cell response | Paclitaxel |  | Dasatinib |  | Dovitinib |  | Pazopanib |  |
| --- | --- | --- | --- | --- | --- | --- | --- | --- | --- |
|  |  | IC<sub>50</sub>(nM) | Hill Slope | IC<sub>50</sub>(nM) | Hill Slope | IC<sub>50</sub>(nM) | Hill Slope | IC<sub>50</sub>(nM) | Hill Slope |
| SiHa | Cervical Cancer Invasion | 47 | -1.00 | 213 | +0.29 ± 0.08 | 89 | -0.44 ± 0.12 | 1,288 | +0.24 ± 0.11 |
| CaSki | | 695 | -0.51 ± 0.12 | 1,248 | -1.00 | 6,363 | +0.51 ± 0.25 | 335 | -0.52 ± 0.21 |
| SiHa | Microvessel Length | 57 | -1.11 ± 0.19 | 139 | -0.86 ± 0.16 | 2 | -1.00 | 944 | -0.50 ± 0.09 |
| CaSki | | 88 | -0.40 ± 0.14 | 94 | -0.75 ± 0.17 | 81 | -1.00 | 126 | -1.97 ± 0.49 |

For cervical cancer invasion and microvessel length, SiHa were more responsive to Paclitaxel compared to Ca Ski. Dasitinib reduced microvessel length in SiHa and Ca Ski equally, yet only Ca Ski were responsive to Dasitinib in terms of cervical cancer invasion. SiHa, in contrast, displayed increasing cervical invasion with increasing Dasitinib dose. Dovitinib reduced the endothelial microvessel length with increasing dose for both SiHa and Ca Ski, with endothelial cells co-cultured with SiHa displaying significantly increased sensitivity to Dovitinib. However, only SiHa were responsive to Dovitinib in terms of cervical cancer invasion whereas Ca Ski displayed increasing cervical cancer invasion with increasing Dovitinib dose. Lastly, Pazopanib also reduced microvessel length with increasing dose for both SiHa and Ca Ski, with endothelial cells co-cultured with Ca Ski displaying significantly increased sensitivity to Pazopanib. Additionally, Pazopanib significantly reduced Ca Ski cervical cancer invasion with increasing dose. However, similar to Dasitinib, the SiHa were resistant to Pazopanib in terms of cervical cancer invasion, with increasing cervical cancer invasion with increased Pazopanib dose. Cervical cancer coverage and endothelial cell coverage were also measured, with endothelial cell coverage following similar trends in terms of sensitivity to those observed using endothelial cell tube length. Similar to cervical cancer invasion, cervical cancer coverage identified SiHa as resistant to Dasitinib and Pazopanib yet did so by identifying a diminished response compared to an increasing response with increased dose. However, cervical cancer coverage did not increase with increasing dose, highlighting the importance of 3D phenotypic screens (**Supplemental Figure 5, Supplemental Table 2**).

## 4. Discussion

Through the use of Design of Experiments, we identified the formulations of two hydrogels in a multilayer multicellular construct of cervical cancer and endothelial cells that simultaneously maximized cancer invasion and endothelial microvessel length. The response surface maps revealed that the ECM protein concentrations and choice of synthetic polymer in both layers significantly affected the cell behavior in both layers, demonstrating the utility of a Design of Experiments approach. Furthermore, by forming the constructs in 96-well plates and using cell phenotype as a read out instead of molecular targets, our tissue engineered model of cervical cancer interfaces with standard high throughput screening methods.

While the majority of high throughput screening is still performed using monolayers of cells on tissue culture plastic, there has been increased interest in using three-dimensional culture models for personalized medicine and drug discovery. This includes bioprinting with natural and synthetic polymers, patient derived organoids with or without biomaterials, and multilayer multicellular constructs.^33–35^ However, hurdles remain to integrate three-dimensional culture models with high throughput high content screening methods, for both molecular target screens and phenotypic screens. Specifically, three of the biggest challenges are engineering platforms that are compatible with liquid handling equipment, quantifying phenotypic response in a high throughput content manner in three dimensions, and mimicking the tissue of interest as closely as possible.^35^ Our platform and analysis methods overcome all of these challenges. By engineering our platforms in a 96-well plate format using IBIDI angiogenesis plates, we are able to interface with automated liquid handling equipment that is standard in high throughput drug screening facilities. This is in contrast to many spheroid cultures that do not interface well with automated liquid handling equipment.^35^ Furthermore, the IBIDI angiogenesis plates are black walled, have a polymer coverslip with extremely low autofluorescence, and enable hydrogels that are 10 μL in volume and 800 μm in height.^36^ This reduces optical light scattering, autofluorescence, and light penetration issues that are prevalent in other larger tissue engineered constructs. Furthermore, by using a high content imaging system that is paired with a live cell analysis system, we are able to automatically observe and calculate cell dynamics in three-dimensions over time. Finally, to mimic the tissue of interest, we chose a multilayer multicellular construct is inspired by the field of human skin equivalent models.^37,38^ Similar to human skin equivalents that we have engineered previously, our model has two distinct layers of hydrogels, two distinct layers of cells, and structural complexity similar to what is observed morphologically in healthy cervical tissue.^30,39,40^ However, the cervical cancer tumor microenvironment has biophysical properties distinct from healthy skin, thus we needed to explore which ECM components to include in our constructs and at what concentrations.

To serve as a screening platform for anti-cancer and anti-angiogenic drugs, *in vitro* models approximating the cervical cancer tumor microenvironment must be able to promote cervical cancer invasion and microvessel formation in the absence of drug. By modulating the constituents of composite hydrogels, we engineered each hydrogel layer to maximize both endothelial cell and cervical cancer cell phenotypes and actions observed during metastasis. All six of the input variables examined (Col1, fibrinogen, and the choice of GelMA or PEGDA in the bottom hydrogel, Col1, fibronectin, and the choice of GelMA or PEGDA in the top hydrogel) had significant effects on the cell responses. Col1 was included in both the bottom hydrogel layer that the endothelial cells are seeded on and the top hydrogel layer that the cervical cancer cells are seeded on as Col1 is an abundant protein surrounding both the cancer cells and the microvessels.^41,42^ We examined Col1 in a range from 0.5 – 2.5 mg/mL as healthy cervical tissue has a collagen concentration of approximately 0.26 mg/mL and we and others have observed that Col1 is upregulated in cervical cancer tissue compared to healthy cervical tissue.^43–45^ We included fibronectin in the top hydrogel layer as fibronectin is increased in cervical cancer compared to healthy cervical tissue and promotes cervical cancer adhesion and invasion.^46,47^ We examined fibronectin in a range from 0.125 – 0.175 mg/mL as 0.01 – 0.10 mg/mL has been used successfully to induce invasion in cervical cancer cells.^48,49^ Additionally, the maximum concentration of fibronectin stock is 1 mg/mL, limiting exploration of higher fibronectin concentrations in our composite hydrogels. In the bottom hydrogel, fibrinogen was included as fibrin is a standard *in vitro* model of endothelial microvessel formation to evaluate angiogenesis and vasculogenesis.^50^ Increasing fibrinogen concentration increases fibril density, fibrin binding sites, and increases substrate stiffness.^51,52^ We examined fibrinogen in a range of 0.5 – 2.5 mg/mL as fibrinogen is found in plasma at a concentration of approximately 3 mg/mL and fibrin hydrogels of 1-2 mg/mL are standard for assays to assess endothelial cell function.^53–55^ Previous studies have eloquently demonstrated that the degree of vasculogenesis of human umbilical cord vein endothelial cells or endothelial progenitor cells can be modulated by tuning the concentration of fibrin and collagen, either alone or in composite hydrogels.^52,56^ Lastly, in addition to modulating the concentrations of the ECM proteins, we evaluated whether GelMA or PEGDA better supported the co-culture model. Our initial screen identified 7% w/v GelMA and 10% w/v PEGDA as the concentrations that best promoted cervical cancer and endothelial cell behavior. These are in range with previous studies that used 2.5 - 10 % w/v GelMA and PEGDA for 3D cancer models.^57–60^

A critical finding of this work is that DOE revealed intimate interactions between all six of the input variables and all four of the output variables. Furthermore, ECM and synthetic polymers in each hydrogel layer significantly influenced the behavior of the cervical cancer cells and the endothelial cells, highlighting the importance of engineering a co-culture construct as a whole instead of optimizing culture conditions for each type of cell and then combining them. This was further demonstrated by the fact that when the cells were cultured on their preferred matrix in the absence of the other cell type, minimal cervical cancer invasion or endothelial microvessel formation was observed. This is most likely due to the fact that in addition to the physical cues from the microenvironment, cancer-endothelial cell crosstalk drives both of these phenotypes in the tumor microenvironment.^61,62^ While communication between cervical cancer cells and endothelial cells has been investigated *in vitro,* all of these studies have been done with either conditioned media, which does not allow for crosstalk, or transwell assays, which do not enable juxtracrine signaling or observation of 3D behavior.^63–65^ Therefore, this work is the first to evaluate the role of cervical cancer-endothelial cell crosstalk on cervical cancer invasion or microvessel formation in an *in vitro* model. The results of our DOE revealed a complex nonlinear relationship between fibrin and collagen concentrations in the bottom hydrogel on both endothelial cell and cervical cancer behavior. These relationships could be obscured if evaluating one factor at a time, as we and others have found that increasing Col1 and fibrinogen concentrations synergistically modulate hMVEC microvessel formation and our construct that maximized all of the cell phenotypes of interest had the maximum concentrations of 2.5 mg/mL Col1 and 2.5 mg/mL fibrinogen in the bottom hydrogel.^66^ The results of our DOE also revealed a complex nonlinear relationship between Col1 and fibronectin in the top hydrogel. There were clear quadratic interactions between Col1 and itself, fibronectin and itself, and linear interactions between Col1 and fibronectin on cervical cancer invasion and coverage. In contrast to the interactions of Col1 and fibrinogen in the bottom hydrogel, the top hydrogel of our construct that maximized all cell phenotypes had concentrations of 1.12 mg/mL Col1 and 0.16 mg/mL fibronectin, which are near the middle of the ranges examined. In comparison to the ECM components which were evaluated as continuous variables, GelMA and PEGDA were evaluated as categorical variables to determine which of the synthetic polymers best supported cancer invasion and endothelial cell microvessel formation. The results of our DOE revealed that both the cervical cancer cells and the endothelial cells had a greater phenotypic response when GelMA was used as the synthetic polymer instead of PEGDA in both hydrogel layers. Both GelMA and PEGDA have been combined with many other natural polymers, enabling flexibility to incorporate ECM components to promote cell function without compromising the diffusion of oxygen, nutrients, or small molecules.^59,67^ Our results complement many other studies that have successfully used GelMA as a 3D culture environment to study cancer invasion, yet is unique in that it identifies the specific microenvironmental conditions of ECM concentrations and synthetic polymers that promote cervical cancer and endothelial cell metastatic behavior simultaneously.

The manipulation of the ECM components and the choice of synthetic polymer altered the mechanical properties of the hydrogels. Human skin is a viscoelastic material with a storage modulus, G’, of approximately 362 Pa and a loss modulus, G”, of 92 Pa.^68^ The hydrogels in our optimized construct are on the same order of magnitude, with storage moduli of 270 and 355 Pa and loss moduli of 20 and 36 Pa. Additionally, the viscoelastic properties of the single polymer hydrogels of collagen, fibrin, GelMA, and PEGDA had similar values to those observed in previous studies.^69–75^ When the ECM proteins are with GelMA the storage modulus is decreased compared to GelMA alone, indicating that the inclusion of the ECM proteins results in a softer hydrogel. Comparing the composition of the top and bottom hydrogels of the optimized construct, the bottom hydrogel has 7% GelMA, has a higher concentration of Col1, and includes fibrin which also forms a hydrogel on its own. In contrast, while the top hydrogel also has 7% GelMA, it has a lower concentration of Col1 and includes fibronectin, which does not form a hydrogel on its own. Thus, the bottom hydrogel having a higher storage and loss modulus compared to the top hydrogel is in line with expected trends in viscoelastic properties as a function of hydrogel composition. Furthermore, the storage and loss moduli of the composite hydrogels are an order of magnitude higher than the hydrogels comprised of the individual ECM proteins alone due to the inclusion of GelMA in the polymer solution. Overall, as expected, changing the composition of the hydrogels influenced material properties. However, including ECM proteins in a GelMA interpenetrating network influences both stiffness and fiber density, thus the differences in cell dynamics as a function of collagen, fibrin, and fibronectin concentrations cannot be attributed to stiffness alone.^76^

To demonstrate that our optimized constructs could integrate with standard drug screening methods, we performed a targeted drug screen of four clinically relevant drugs on two cervical cancer cell lines. Treating the cervical cancer cells with Paclitaxel, a chemotherapeutic agent commonly used in cervical patients,^77^ resulted in more shallow curves and robust phenotypic responses compared to the other drugs assessed. The IC_50_ values of 47 nM and 695 nM for SiHa and Ca Ski invasion, respectively, demonstrate that Ca Ski had limited responsiveness to Paclitaxel. This trend of SiHa exhibiting increased sensitivity to Paclitaxel was also observed for cervical cancer coverage. This is in contrast with studies that examined Paclitaxel sensitivity in Ca Ski and SiHa using 2D tissue culture plastic and found that SiHa had a higher IC_50_ value for Paclitaxel compared to Ca Ski.^78,79^ However, those studies used cell viability of cervical cancer cells in monoculture to calculate the IC_50_ values, whereas we evaluated the phenotypic responses of cervical cancer cells in co-culture, highlighting the fact that cancer invasion does not always follow the same trends as cell death. Furthermore, cancer cells in 3D have significantly increased chemoresistance compared to those in 2D, as do cells in co-culture compared to those in monoculture.^80,81^ Additionally, similar to previous studies, Paclitaxel induced an anti-angiogenic response and the endothelial cells were more sensitive to Paclitaxel compared to the cervical cancer cells.^82^ Dasitinib, Dovitinib, and Pazopanib are all tyrosine kinase inhibitors that are used clinically as anti-angiogenic agents.^82–86^ All three of these drugs induced an antiangiogenic response in terms of reduced microvessel length and reduced endothelial cell coverage with increasing dose intensity. However, all three of these drugs also resulted in increasing cervical cancer invasion for one of cervical cancer cell lines examined. Specifically, Dasitinib and Pazopanib induced increasing cervical cancer invasion with increasing dose intensity for SiHa and Dovitinib induced increasing cervical cancer invasion for CaSki. Furthermore, even though Ca Ski was responsive to Dasitinib and Pazopanib and SiHa were responsive to Dovitinib, the IC_50_ values were over 1 μM and should be treated with caution. Taken together, these data indicate that these three drugs alone would most likely not be effective treatment strategies for either of these cervical cancer patients. This is in line with numerous other studies that have found that antiangiogenic drugs alone are not sufficient to inhibit metastasis and to be clinically effective they should be co-delivered with anti-tumor agents such as Paclitaxel.^83,85^ Our screening platform could be used for future studies that investigate combinatorial treatment strategies of these and other drug regimens.

## 5. Conclusion

We developed and validated a 3D *in vitro* model of the cervical cancer tumor microenvironment that is compatible with liquid handling equipment used in standard high throughput drug screening methods, quantifies phenotypic response of both cervical cancer and endothelial cells in three dimensions over time, and has greater phenotypic responses compared to the current gold standard of Matrigel. By using DOE, we identified the specific microenvironmental conditions of ECM concentrations and synthetic polymers that maximized cervical cancer and endothelial cell metastatic behaviors simultaneously. In conclusion, we developed a phenotypic drug screening platform of cervical cancer that captures cell behavior present in the cervical cancer tumor microenvironment, captures patient to patient variability, and integrates with standard high throughput high content drug screening methods. This platform can be used for future studies to screen large compound libraries for drug discovery and precision oncology.^87,88^

---

**Supplemental Table 1.** Input variable combinations examined in D-optimal DOE design.

| Exp No | Exp Name | Run Order | Incl/Excl | Collagen Top | Fibronectin Top | Fibrinogen Bottom | Collagen Bottom | Synthetic Polymer Top | Synthetic Polymer Bottom |
| --- | --- | --- | --- | --- | --- | --- | --- | --- | --- |
| 1 | N1 | 11 | Incl | 0.5 | 0.175 | 0.5 | 0.5 | PEGDA | PEGDA |
| 2 | N2 | 23 | Incl | 0.5 | 0.125 | 2.5 | 0.5 | PEGDA | PEGDA |
| 3 | N3 | 12 | Incl | 2.5 | 0.175 | 2.5 | 0.5 | PEGDA | PEGDA |
| 4 | N4 | 32 | Incl | 0.5 | 0.175 | 2.5 | 2.5 | PEGDA | PEGDA |
| 5 | N5 | 7 | Incl | 2.5 | 0.175 | 1.16667 | 2.5 | PEGDA | PEGDA |
| 6 | N6 | 1 | Incl | 2.5 | 0.141667 | 0.5 | 0.5 | PEGDA | PEGDA |
| 7 | N7 | 20 | Incl | 1.83333 | 0.125 | 0.5 | 2.5 | PEGDA | PEGDA |
| 8 | N8 | 34 | Incl | 0.5 | 0.175 | 2.5 | 0.5 | GELMA | PEGDA |
| 9 | N9 | 10 | Incl | 0.5 | 0.125 | 2.5 | 2.5 | GELMA | PEGDA |
| 10 | N10 | 6 | Incl | 0.5 | 0.125 | 0.5 | 1.16667 | GELMA | PEGDA |
| 11 | N11 | 21 | Incl | 2.5 | 0.125 | 2.5 | 1.16667 | GELMA | PEGDA |
| 12 | N12 | 2 | Incl | 2.5 | 0.125 | 1.16667 | 0.5 | GELMA | PEGDA |
| 13 | N13 | 30 | Incl | 2.5 | 0.175 | 0.5 | 1.16667 | GELMA | PEGDA |
| 14 | N14 | 18 | Incl | 2.5 | 0.158333 | 2.5 | 2.5 | GELMA | PEGDA |
| 15 | N15 | 13 | Incl | 1.16667 | 0.175 | 0.5 | 2.5 | GELMA | PEGDA |
| 16 | N16 | 8 | Incl | 0.5 | 0.175 | 2.5 | 0.5 | PEGDA | GELMA |
| 17 | N17 | 19 | Incl | 0.5 | 0.125 | 0.5 | 2.5 | PEGDA | GELMA |
| 18 | N18 | 5 | Incl | 2.5 | 0.175 | 0.5 | 2.5 | PEGDA | GELMA |
| 19 | N19 | 4 | Incl | 2.5 | 0.125 | 2.5 | 2.5 | PEGDA | GELMA |
| 20 | N20 | 31 | Incl | 2.5 | 0.125 | 1.83333 | 0.5 | PEGDA | GELMA |
| 21 | N21 | 24 | Incl | 1.83333 | 0.125 | 0.5 | 0.5 | PEGDA | GELMA |
| 22 | N22 | 17 | Incl | 1.83333 | 0.175 | 2.5 | 2.5 | PEGDA | GELMA |
| 23 | N23 | 26 | Incl | 1.5 | 0.15 | 1.5 | 1.5 | PEGDA | GELMA |
| 24 | N24 | 14 | Incl | 2.5 | 0.125 | 0.5 | 2.5 | GELMA | GELMA |
| 25 | N25 | 9 | Incl | 0.5 | 0.175 | 0.5 | 1.83333 | GELMA | GELMA |
| 26 | N26 | 15 | Incl | 0.5 | 0.141667 | 0.5 | 0.5 | GELMA | GELMA |
| 27 | N27 | 3 | Incl | 0.5 | 0.158333 | 2.5 | 2.5 | GELMA | GELMA |
| 28 | N28 | 16 | Incl | 2.5 | 0.175 | 2.5 | 1.83333 | GELMA | GELMA |
| 29 | N29 | 28 | Incl | 2.5 | 0.175 | 1.83333 | 2.5 | GELMA | GELMA |
| 30 | N30 | 33 | Incl | 2.5 | 0.158333 | 2.5 | 0.5 | GELMA | GELMA |
| 31 | N31 | 35 | Incl | 1.16667 | 0.125 | 2.5 | 0.5 | GELMA | GELMA |
| 32 | N32 | 29 | Incl | 1.16667 | 0.175 | 0.5 | 0.5 | GELMA | GELMA |
| 33 | N33 | 27 | Incl | 1.5 | 0.15 | 1.5 | 1.5 | GELMA | GELMA |
| 34 | N34 | 22 | Incl | 1.5 | 0.15 | 1.5 | 1.5 | GELMA | GELMA |
| 35 | N35 | 25 | Incl | 1.5 | 0.15 | 1.5 | 1.5 | GELMA | GELMA |

**Supplemental Table 2.** Comparison of the inhibitory effect of different drugs in multilayer multicellular *in vitro* model for cervical cancer. IC_50_ and Hill Slope values for endothelial cell coverage and cervical cancer coverage. Values represent mean ± SEM, N=3.

| Cell line | Inhibitory Agent<br>Cell response | Paclitaxel |  | Dasatinib |  | Dovitinib |  | Pazopanib |  |
| --- | --- | --- | --- | --- | --- | --- | --- | --- | --- |
|  |  | IC<sub>50</sub>(nM) | Hill Slope | IC<sub>50</sub>(nM) | Hill Slope | IC<sub>50</sub>(nM) | Hill Slope | IC<sub>50</sub>(nM) | Hill Slope |
| SiHa | Cervical cancer coverage | 90 | -0.36 ± 0.07 | 211 | -0.36 ± 0.12 | 1 | -1.00 | 1 | -0.14 ± 0.14 |
| CaSki |  | 453 | -0.32 ± 0.09 | 598 | -0.46 ± 0.12 | 102 | -1.00 | 315 | -1.00 |
| SiHa | Endothelial cell coverage | 110 | -0.73 ± 0.11 | 413 | -0.24 ± 0.09 | 28 | -0.29 ± 0.03 | 359 | -0.64 ± 0.21 |
| CaSki |  | 42 | -0.38 ± 0.11 | 198 | -0.70 ± 0.26 | 139 | -0.34 ± 0.14 | 317 | -1.00 |

## Notes

### Competing Interest Statement

The authors have declared no competing interest.

