## Supplemental Figures for "Engineering High Throughput Screening Platforms of Cervical Cancer"

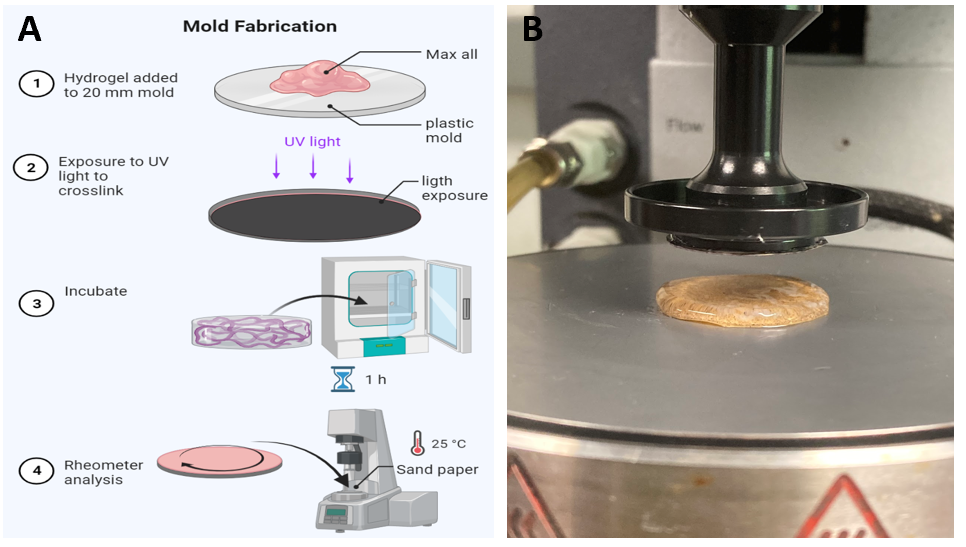

**Figure S1.** Hydrogels characterization **(A)** Schematic representation of the mold fabrication for the rheometer test **(B)** Representative image of the hydrogel combination in the rheometer before compression (Max all bottom layer combination). Hydrogels were photo-cross-linked (λ = 365 nm) and incubated at 37 $℃$ for 1-hour prior rheometer test. Samples were placed on a 20mm standard steel parallel top plate with a bottom standard Peltier plate. Frequency sweeps were conducted from 0.1 to 100 rad/s within the linear viscoelastic (LVE) region at a constant strain amplitude of 5%. Due to the tendency for the gels to slip, 150 grit (120 µm particle size) sandpaper was adhered to both the top and bottom fixtures

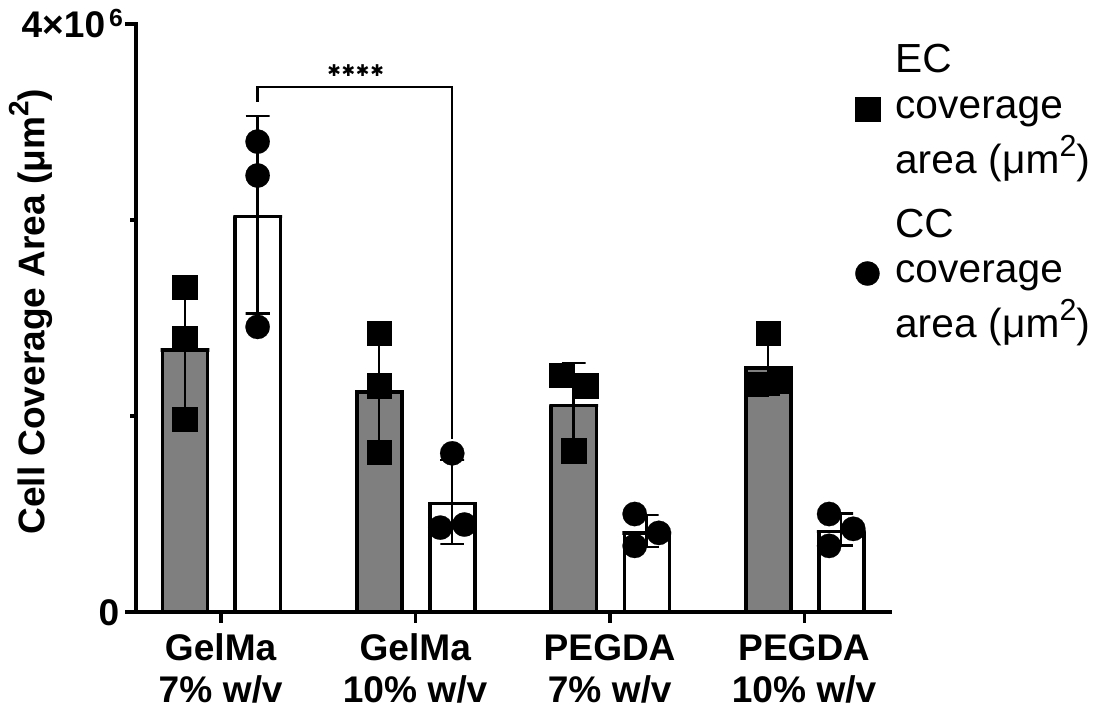

**Figure S2.** Cell coverage area comparison between the synthetic polymers GelMA and PEGDA. Endothelial cell (EC) Cervical cancer cells (CC). * p <0.03, ** p<0.0021, *** p <0.0002 compared with the mean of each group. Two-way ANOVA with Tukey post-test Data represents the mean ± SD (n =3). Hydrogels were photo-cross-linked (λ = 365 nm) and incubated at 37 $℃$ for 48h. Data represents 24h after cultured.

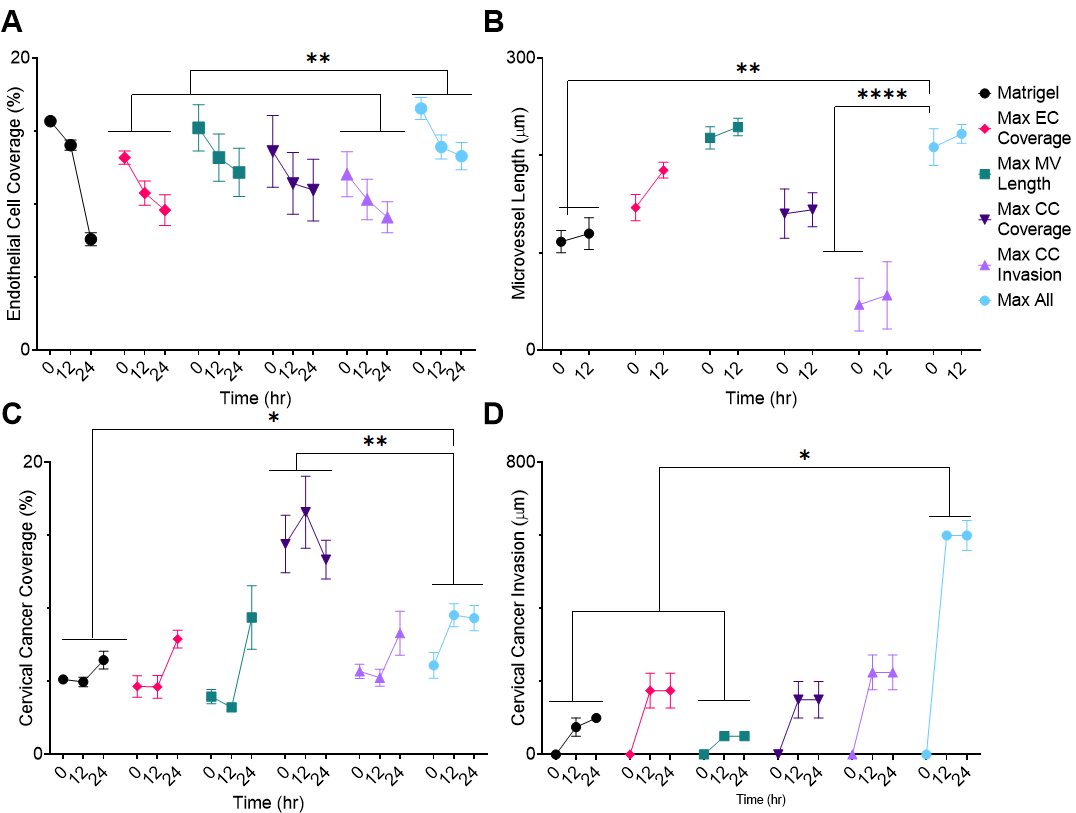

**Figure S3.** Validation of a DOE optimization model in cervical cancer cells (**A – D**) hMVECs and SiHa were co-cultured using the optimized hydrogel combinations to maximize each cell response. Cells were cultured for 48hr. (**A**) endothelial cells coverage (**B**) microvessel length (**C**) cervical cancer coverage (**D**) cervical cancer invasion. (CC) cervical cancer cells, (EC) endothelial cells. * p <0.03, ** p<0.0021, *** p <0.0002 compared with the mean of each group. Two-way ANOVA with Tukey post-test. Data represent the mean ± SEM (n =4).

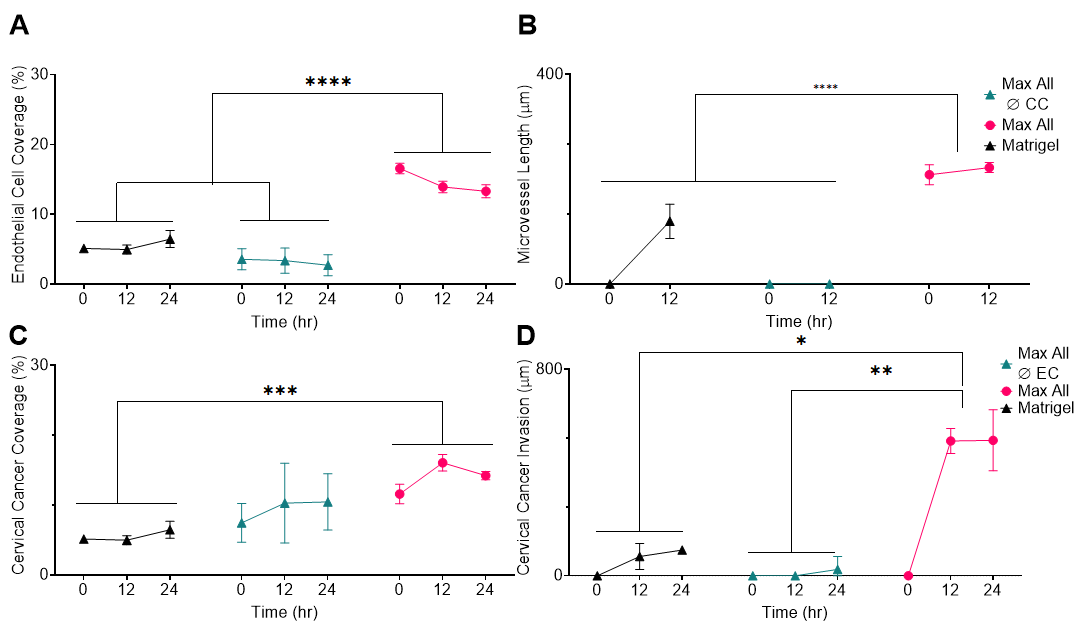

**Figure S4.** Importance of 3D co-culture model **(A – D)** hMVECs and SiHa were cultured

alone and co-cultured using the optimized hydrogel (max all). Data showed after 24h of cultured. **(A)** Endothelial cell coverage **(B)** Microvessel length **(C)** Cervical cancer coverage **(D)** Cervical cancer invasion. (CC) cervical cancer cells, (EC) endothelial cells. * p <0.03, ** p<0.0021, *** p <0.0002 compared with the mean of each group. Two-way ANOVA with Tukey post-test Data represents the mean ± SEM (n =4

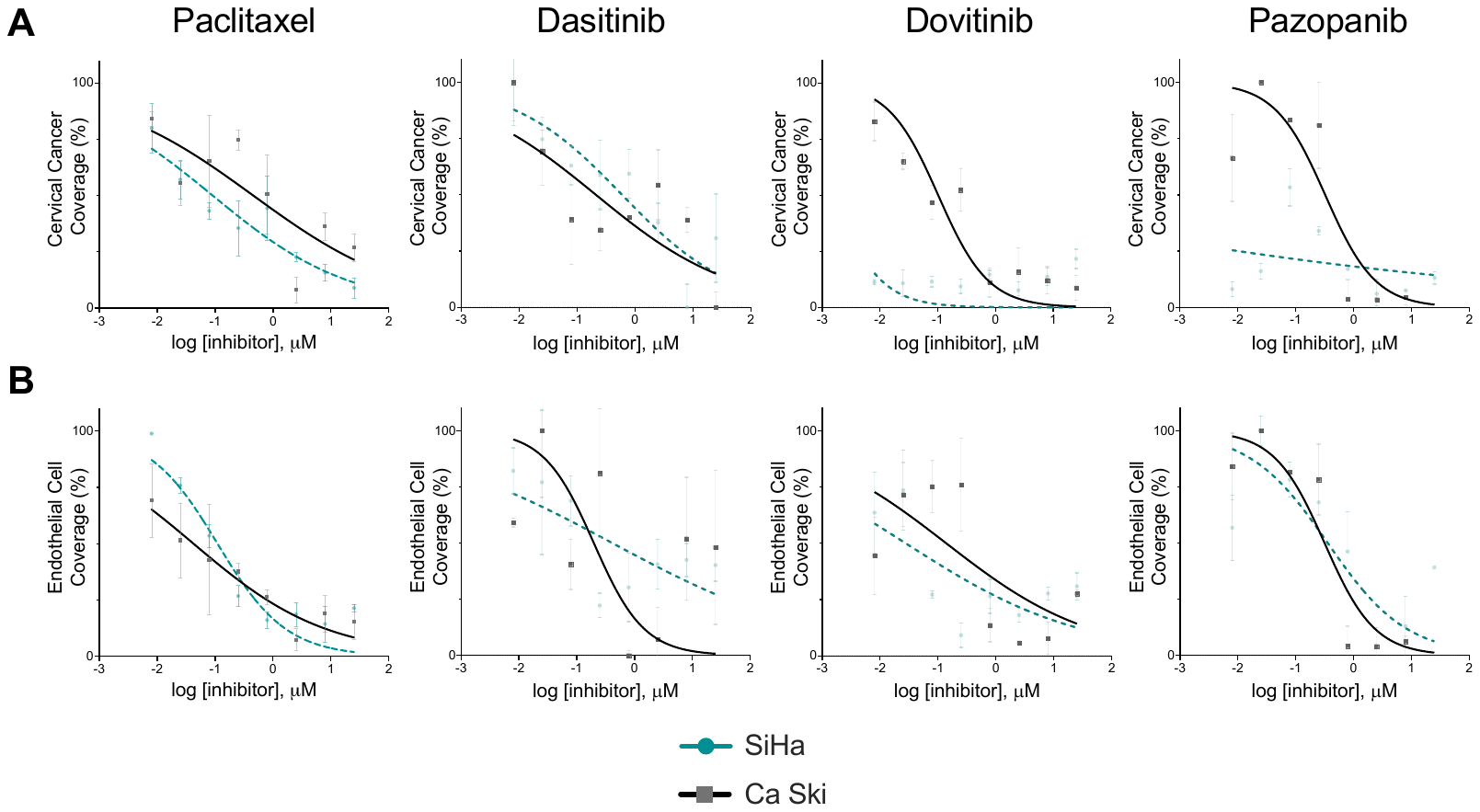

**Figure S5.** *In vitro* growth-inhibitory effects of the inhibitory drugs on human microvascular endothelial cells (hMVECs) and human cervical cancer cell lines (SiHa, Ca Ski). Cells were seeded in the construct, cultured for 24 hours, then treated with 0.008 – 25 µM of drug for 24 hours, at which point cell response was evaluated. Each cell response is normalized to the average of the values observed in the absence of drug. **(A)** Cervical cancer coverage **(B)** Endothelial coverage. Data represent the mean ± SD (n = 3).
